# WormTracer: A precise method for worm posture analysis using temporal continuity

**DOI:** 10.1101/2023.12.11.571048

**Authors:** Koyo Kuze, Ukyo T. Tazawa, Karin Suwazono, Yu Toyoshima, Yuichi Iino

## Abstract

This study introduces WormTracer, a novel algorithm designed to accurately quantify temporal evolution of worm postures. Unlike conventional methods that analyze individual images separately, WormTracer estimates worm centerlines within a sequence of images concurrently. This process enables the resolution of complex postures that are difficult to assess when treated as isolated images. The centerlines obtained through WormTracer exhibit higher accuracy compared to those acquired using conventional methods. By applying principal component analysis to the centerlines obtained by WormTracer, we successfully generated new eigenworms, a basic set of postures, that enables a more precise representation of worm postures than existing eigenworms.

**Author summary:** *C. elegans* is a valuable model organism for comprehensive understanding of genes, neurons and behavior. Quantification of behavior is essential for clarifying these relationships, and posture information plays a crucial role in the analyses. However, accurately quantifying the posture of *C. elegans* from video images of worms is challenging, and while various methods have been developed to date, they have their own limitations.

In this study, we developed an analytical tool called WormTracer, which can obtain worm centerlines more accurately than conventional methods, even when worms assume complex postures. Using this tool, we successfully obtained new eigenworms, basis postures of a worm, that can more accurately reproduce various postures than conventional eigenworms. WormTracer and the new eigenworms will be valuable assets for future quantitative studies on worm locomotion and sensorimotor behaviors.

## Introduction

*C. elegans* is an animal with a fully described body plan where all 959 somatic cells have been named. Furthermore, its nervous system consists of only 302 neurons, and electron microscopy has revealed all the connections between these neurons (connectomes) [1]. It is a valuable model organism for studying genetics and neuroscience. In these fields, numerous studies are being conducted including those on the effects of genetic variation on behavior [2][3] and relationship between neural activity and behavior [4][5][6][7].

To conduct such studies, it is necessary to accurately quantify the behaviors of worms. Since worms have rod-like body shapes, their postures can be represented by a single curved line (centerline) passing through the center of the worm. By approximating the centerline of a worm by 100 linear segments and representing the shape by a vector of angles of each segment, and performing principal component analysis of the vectors, around five principal components, or eigenworms, were found to be sufficient for effectively representing the worm’s postures when used as basis vectors [8].

In many cases, obtaining the centerline of a worm from an image can be easily achieved by combining binarization and thinning [9]. The thinning method is a technique in which, by calculating the configuration of neighboring pixels, the values of pixels lying on edges of a blob is set to zero, gradually thinning the body region to a single line.

However, worms sometimes assume complex postures, such as curled up and crossed over postures. These complex postures are exhibited during fundamental behaviors during locomotion, such as omega turns (movements in which the worm curls up to make the head and tail touch each other, as a trigger for changing the locomotion direction) [10], foraging [11][12], chemotaxis [13], and avoidance responses to nociceptive stimuli [14]. Also, there are certain mutations that make these movements more frequent [3]. However, identifying the centerline using simple methods like thinning becomes difficult when worms adopt such complex postures. In this study, postures for which the centerline cannot be obtained by thinning are referred to as complex postures, while postures for which the centerline can be easily obtained through thinning method are referred to as simple postures.

Various methods have been developed to analyze worms in complex postures, including machine learning using convolutional neural network (CNN) [15], optimization by searching through a low-dimensional space spanned by eigenworms [16], estimating the centerline from image features such as edges [3], and evaluation using multiple criteria, such as previous and following frames and colors, to deal with overlaps between worms [17]. However, methods that utilize complex models like [15][16] can be difficult to adapt to user-specific data. Additionally, approaches that rely on worm texture or luminance value distributions to estimate the posture [15][16][17][18] can be affected by image acquisition conditions. Using information such as the texture of the worm or difference in brightness, for example between the tip and the center of the worm, is not effective under image acquisition conditions where the texture of the worm is not clearly visible or temporally variable. In such cases, using these grayscale pixel values to estimate posture can be counterproductive.

This study proposes a method for estimating the posture of worms using only binarized video, which can be applied to a wide variety of image acquisition conditions. The method addresses the issue of multiple sub-optimal solutions by leveraging information from the postures before and after the occurrence of complex postures. Additionally, the use of GPU-based optimization with the stochastic gradient method allows for rapid estimation.

In addition, the head of *C. elegans* is known to be under sophisticated motor control which is largely different from that of the body. In studies that focus on fine head movements, such as neck swiveling [19][20][21][22], it is necessary to obtain the posture of the worm’s ends with a particularly high precision. However, existing methods have encountered significant challenges in obtaining the posture of the worm ends, limiting their ability to analyze fine motions. In this study, by adjusting the shape of a reconstructed worm during optimization, a centerline could be obtained that closely approximates the actual posture of the whole body, including the ends of the worm.

The commonly used eigenworms are based on principal components obtained from centerline data that do not include complex postures. Therefore, its representation capability of complex postures may be limited. However, since our method allows for the estimation of worm centerlines with high accuracy even in complex postures, we succeeded in computing eigenworms with an improved representation power.

## Results

### Overview of WormTracer

To obtain centerlines of a worm, WormTracer uses only binary images as an input. Firstly, at step 0, the candidate centerlines are obtained by the thinning algorithm of the binary images for all frames (Fig 1). The worm images are reconstructed based on the obtained centerlines, and the discrepancy between the reconstructed image and the actual binary image is used to determine whether the centerlines are successfully obtained at this step. A centerline is successfully obtained for most cases when the worm assumes a simple posture.

**Fig 1.**
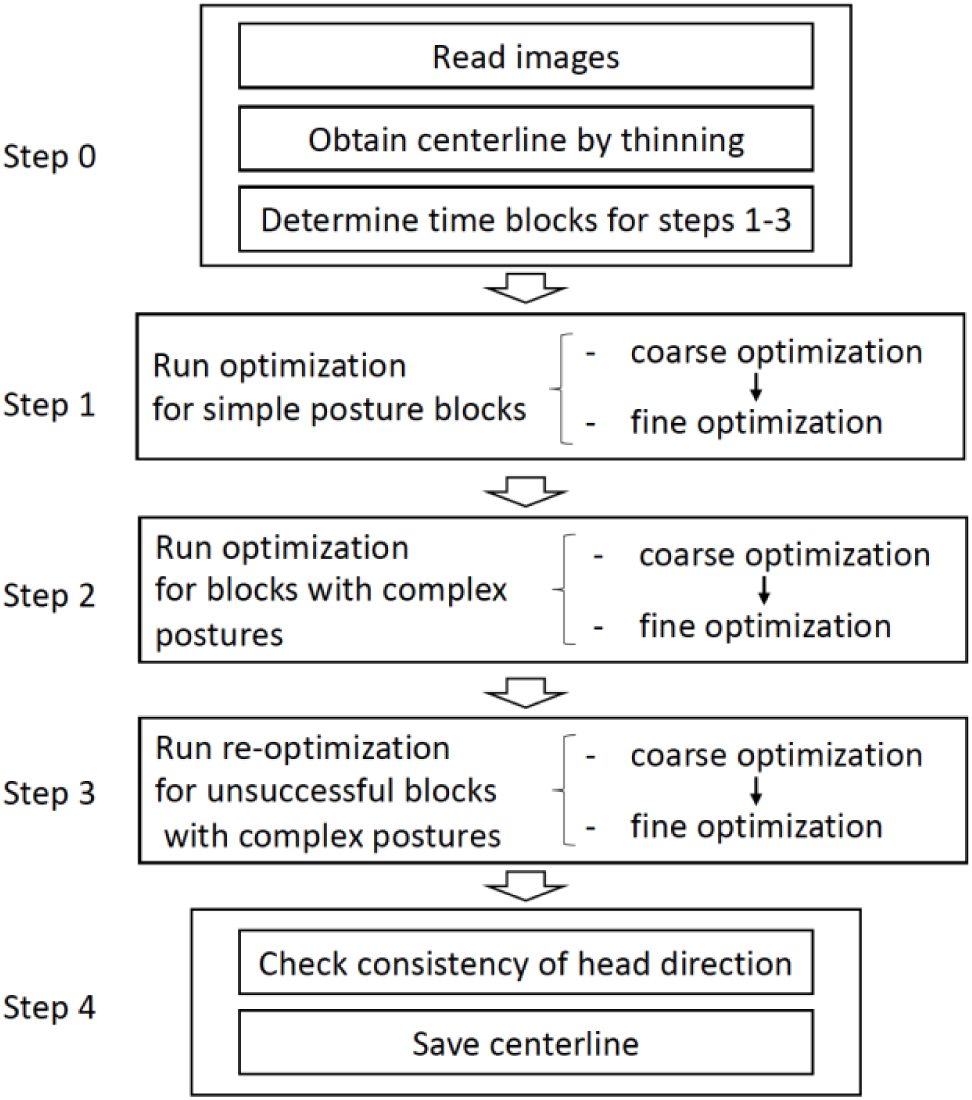
Overview of WormTracer.

This information is then utilized to identify blocks of frames where the centerlines were not accurately obtained, and the whole time series is divided into blocks for gradient-based optimization at following steps, so that each cluster without accurate centerline information is fully included in a block (Fig 2).

**Fig 2.**
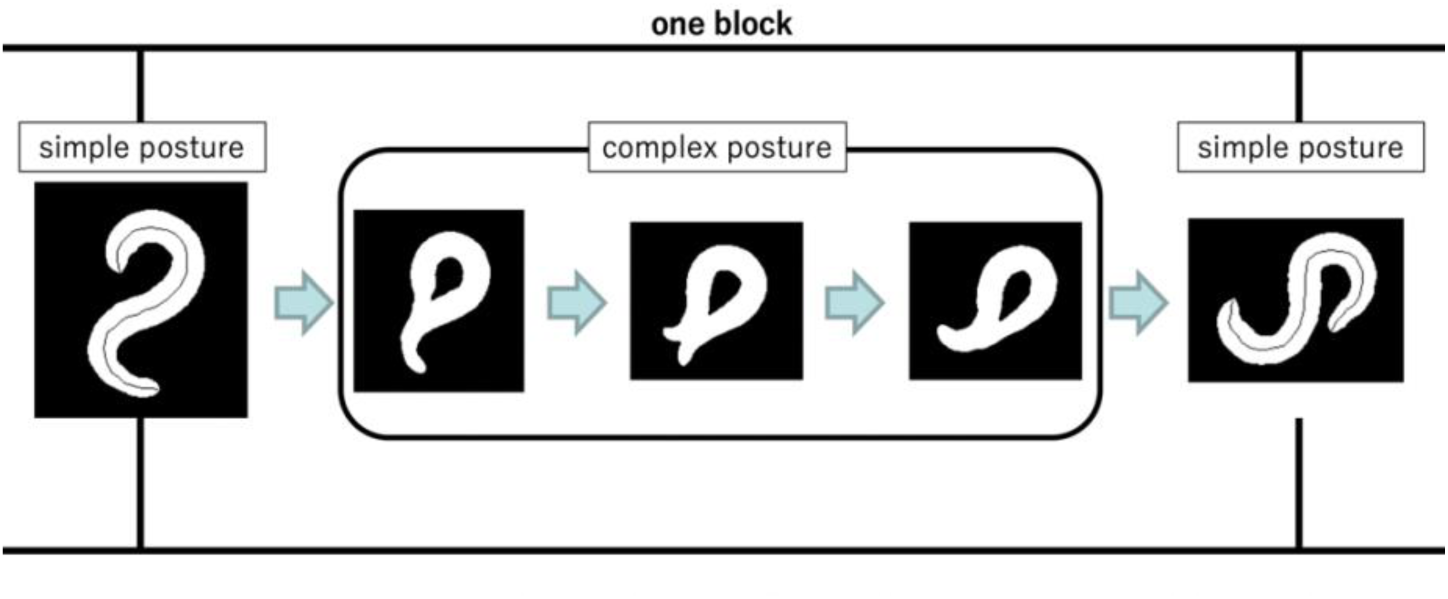
blocks for gradient-based optimization. Each block includes one or zero sequential complex posture images. The beginning and ending images are simple postures.

For each block, exact centerlines are estimated using a stochastic gradient method for optimization. The optimization process is divided into three main steps:

Step 1: Step 1 is performed for blocks with only simple postures where the thinning algorithm was successful. For these blocks, the step 0 centerlines are further refined by gradient optimization.

Step 2: In the second step, optimization is performed for blocks including complex postures that the thinning algorithm did not provide useful candidate centerlines. In this case, as described above, the ends of the blocks are frames with centerline information obtained by thinning, and this information is used to estimate the centerline of the complex postures at the middle of the blocks.

Step 3: In the third step, if the centerline estimated in step 2 is deemed incorrect, the optimization is repeated with different initial values for gradient optimization.

Each of the optimization steps 1 to 3 is in fact divided into the first half and the second half. During the first half, the posture of the worm is coarsely estimated, and during the second half, the details of the centerlines are fine-tuned.

Finally, at step 4, the head and tail of the centerlines are checked to ensure that they are not reversed at the junctions of the blocks, and the centerlines are saved after estimating the orientation of the head throughout the time series (See Materials and Methods for more details).

### Processing time and required computing resources

We recommend a GPU-enabled environment for running WormTracer; without GPU computation, the processing time will be very long. For example, in Google Colaboratory, a K20 GPU can complete the processing of 4200 120x120 images in 120 min, and a T4 GPU can complete the processing in 20 min.

### Comparison with existing methods

Results obtained with previously proposed methods, Wormpose (WP) [15] and EigenWormTracker (EWT) [16], were used for comparison with WormTracer (WT). To conduct the comparison, we applied each of the three methods to the sample data provided in respective reports and compared the results with manually generated ground truth centerlines. Scores were calculated by averaging the distance of each segment junction point from that of the ground truth centerline.

By comparing the error values for each frame, it was found that there are frames where all three methods had weakness, while there were also frames where one or two of the methods performed particularly better compared to other methods (Fig 3a-c). Averaging the results across all frames revealed that the centerlines obtained with WormTracer exhibited the highest accuracy across all types of images (Fig 3d).

**Fig 3.**
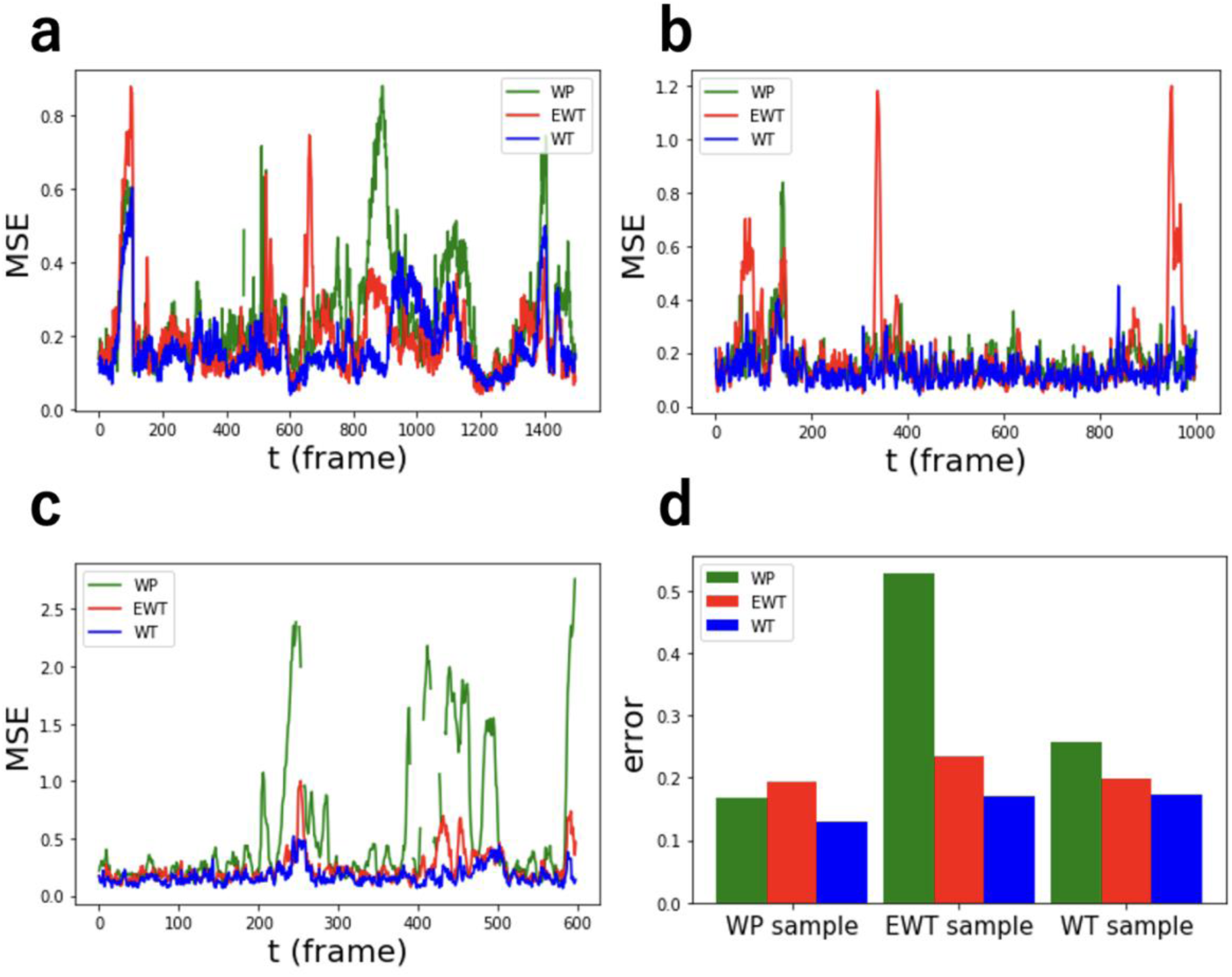
Centerline errors obtained for the sample data of the three methods, displayed in the time direction. a: Mean squared error (y axis) of estimated centerline for a WT sample data for each time point (x axis) b: Mean squared error (y axis) of estimated centerline for a WP sample data for each time point (x axis) c: Mean squared error (y axis) of estimated centerline for a EWT sample data for each time point (x axis) d: Bar graph of centerline errors obtained for the sample data of the three methods, averaged over all frames for the three methods Color corresponds to each estimation method. For all three samples, the result using WT has the smallest error.

To investigate the strengths and weaknesses of the three methods, we compared the centerlines’ actual placements for different postures. In the case of WormTracer sample data, all three methods had higher error values around the frame *t*=100 (Fig 3a). During this period, the worm curled up, and the head appeared to be stuck in the body (Fig 4a), resulting in all methods producing centerlines with elongated heads compared to their actual positions. In the WP sample data, EWT analysis did not perform well in some instances (Fig 3b). For example, at *t*=940, despite the worm being in a simple posture, the centerline was misaligned by EWT (Fig 4b). This observation suggests that the postures represented in low-dimensional space by eigenworms, which EWT was based on, might have only limited representation power. In the EWT sample data, WP analysis did not perform well overall (Fig 3c). For instance, at *t*=450, the rounded shape did not correctly match the body’s actual shape (Fig 4c). WP heavily relies on grayscale patterns, but in the case of the EWT sample data, these functionalities might not have been effective due to the lack of clear patterns in this image set.

**Fig 4.**
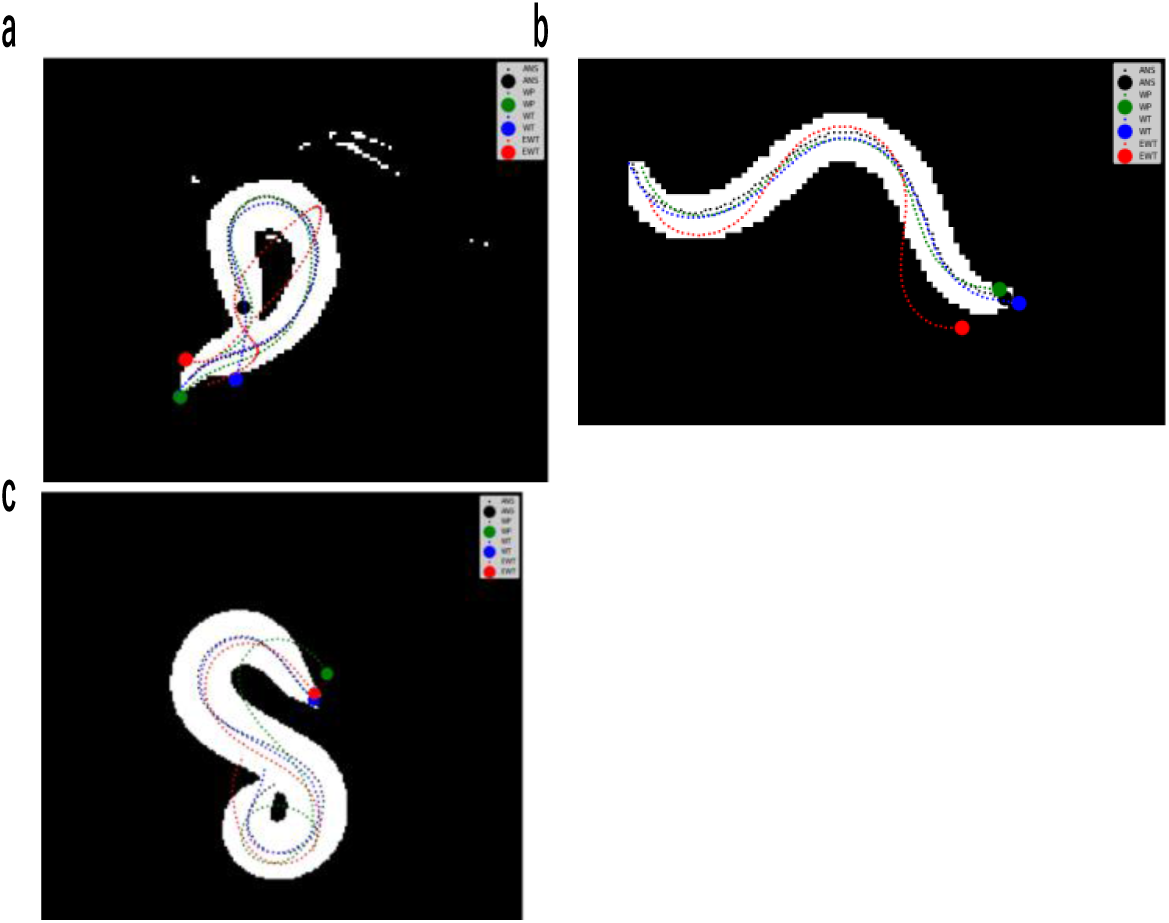
Centerlines obtained by the three methods superimposed on a binary input image of the worm. a: WormTracer sample data. The worm is curled up, and the positions of the tip obtained by all methods are incorrect. b: Sample image from WP; centerline estimation by EWT is significantly displaced. c: Sample image from EWT; centerline estimation by WP is significantly displaced.

There were no frames in any of the sample data where the results of WT were obviously inferior compared to the other methods.

### Validation with different types of image data

To further evaluate the dependence of the performance of our proposed method on the nature of the image data, different types of data were prepared. First, WormTracer was run on 5 fps video instead of 66 fps, and it successfully acquired the appropriate centerlines, just like the 66 fps data (Fig 5). This indicates that the method is robust to variations in the frame rate of the video.

**Fig 5.**
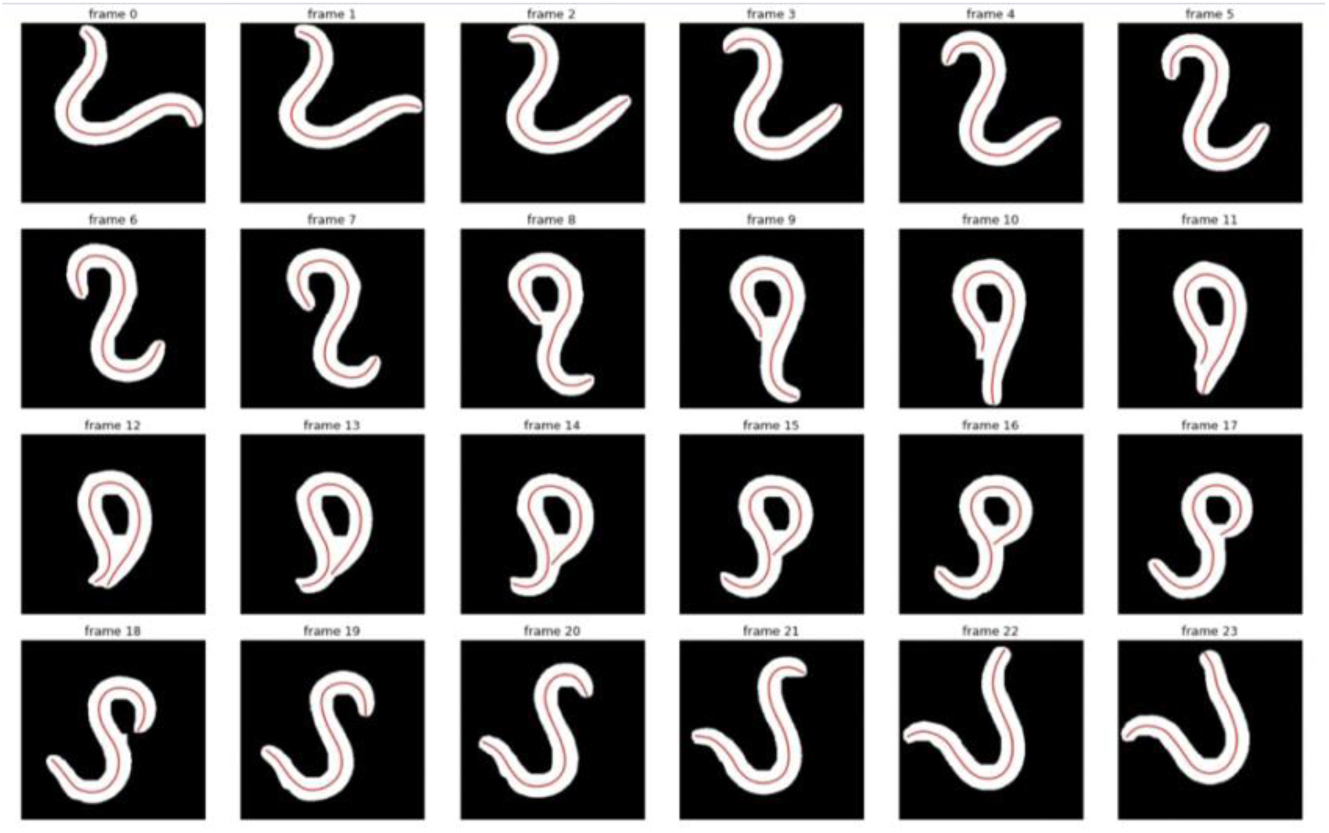
Result of running WormTracer on images taken at 5 fps. The binarized images of the worm (white) and the center line output by WormTracer (red) are shown.

Next, WormTracer was applied to images of the *dpy* (dumpy) mutant, which has a stubby shape compared to wild-type worms. The hyperparameters used for the optimization were exactly the same as those used for wild-type worms. As a result, WT was found to be applicable to such images (Fig 6).

**Fig 6.**
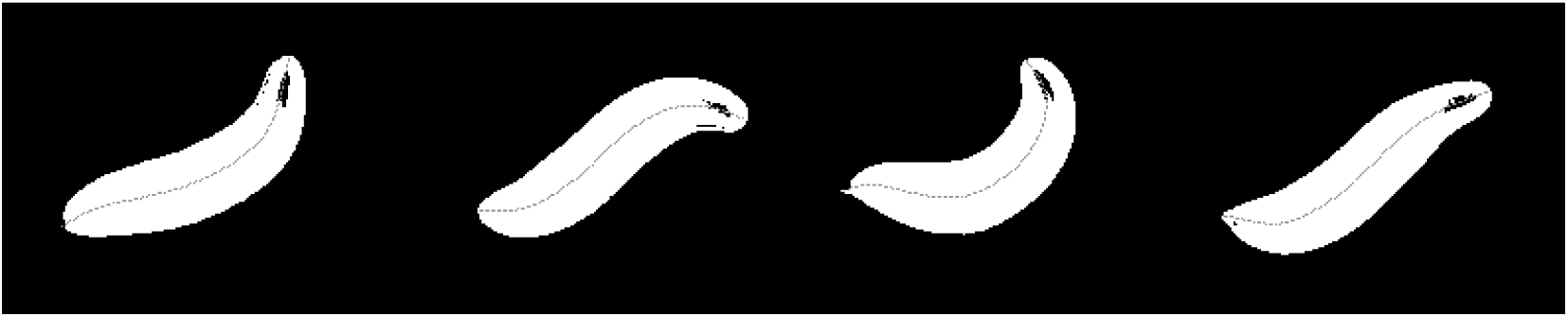
Centerlines were obtained using WormTracer for the images of the dpy mutant.

### Creation of new eigenworms and evaluation of the expressive capacity

Eigenworms have been proposed based on the idea that the posture of worms is formed by a weighted sum of a small number of basis patterns, and commonly used eigenworms are principal components of the centerlines obtained from images that do not include complex postures (Fig 7a) [23]. Hence, it should be possible to derive a more general eigenworm by performing a principal component analysis on the centerlines obtained by WormTracer applied to images that included complex postures. Accordingly, we obtained a new set of eigenworms from centerlines obtained by applying WormTracer to images from the Open Worm Movement Database [24], which includes large-scale worm behavior images (Fig 7b).

**Fig 7a.**
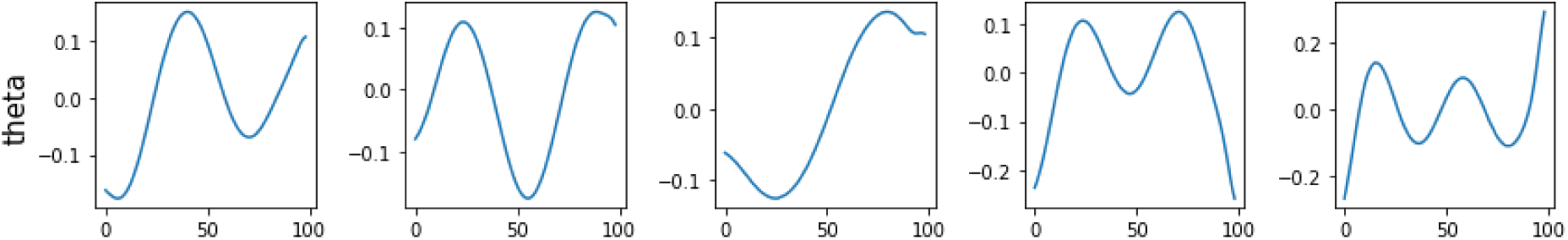
Previously reported eigenworm. From left to right: first principal component, second principal component, and so on. x axis shows segment numbers and y axis shows the angle of segment lines.

**Fig 7b.**
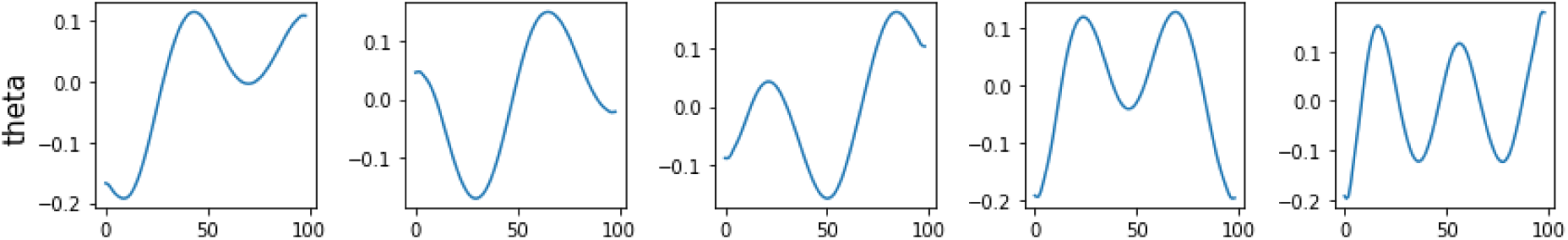
Newly obtained eigenworm. From left to right: first principal component, second principal component, and so on. x axis shows segment numbers and y axis shows the angle of segment lines.

To examine the representation power of these eigenworms, we reconstructed centerline data obtained by applying WT to each of the sample videos of WT, EWT, and WP, using both the old and new eigenworms. Since these sample images frequently assume complex postures, it is crucial to assess whether the reconstructions perform well for such postures. The errors were calculated by comparing the theta values (angles of centerline segments) reconstructed using each eigenworm with the theta values obtained by WT. It was found that the new eigenworm represented the centerline more accurately for all sample data (Fig 8). Moreover, the errors along the body axis showed that the new eigenworm was particularly more precise at both ends compared to the old eigenworm (Fig 8).

**Fig 8.**
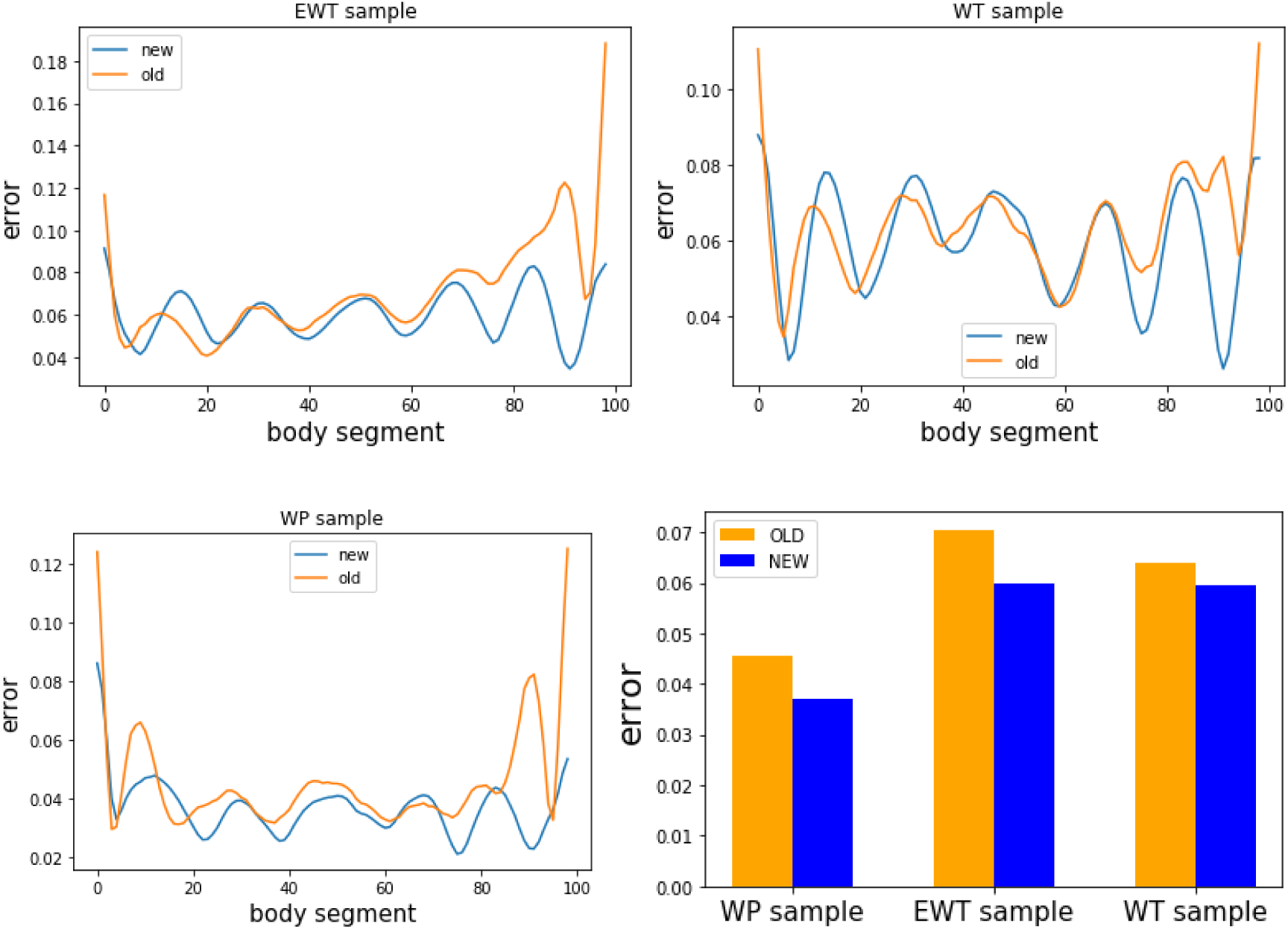
The difference between the angle reconstructed using the existing eigenworm (orange) and the new eigenworm (blue), and the angle obtained by WormTracer. Overall, the new eigenworm is better and more accurate, especially for both ends of the centerline.

The reconstructed centerlines using the old and new eigenworms were superimposed along with the centerlines obtained by WT (Fig 9). The centerlines reconstructed with the old eigenworm were found to have misalignments at both ends in some frames.

**Fig 9.**
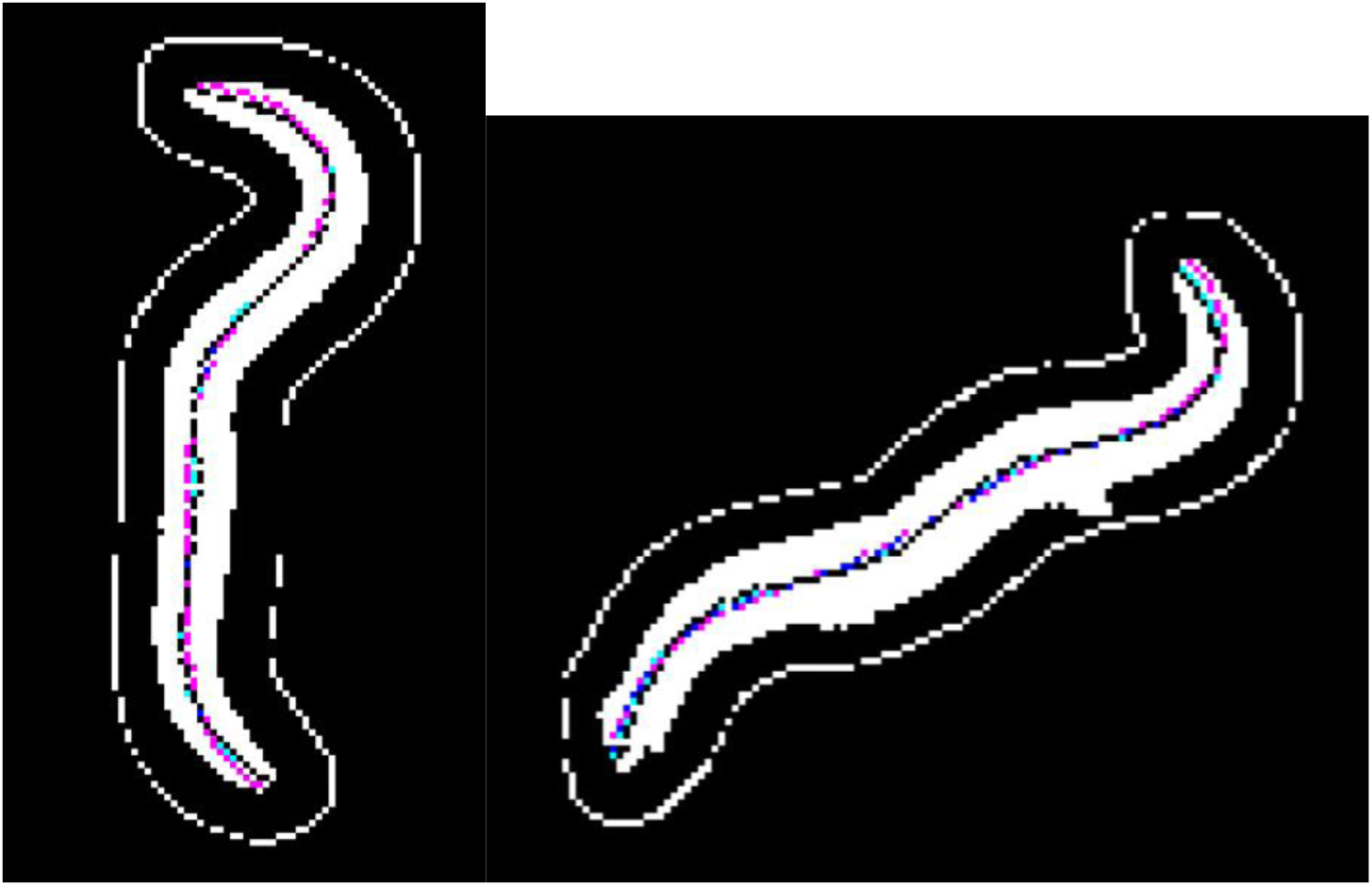
Centerlines obtained by WormTracer (black), centerlines reconstructed using existing eigenworms (red) and those with the new eigenworms (blue) are shown superimposed on binarized input images.

## Discussion

WormTracer can easily and quickly obtain the centerlines of a worm once users prepare binarized images from moving images of the worm. In terms of accuracy, this method can obtain the correct centerlines even for images where conventional methods fail to do so. It is also capable of obtaining the correct centerlines in difficult postures where multiple candidates for the centerlines are found.

One advantage of using binarized images is that they are not easily affected by imaging conditions. If the image is taken under conditions that do not reflect the body texture, the texture information cannot be used. In a setup where illumination is unevenly directed, the grayscale images will give incorrect information if the brightness varies depending on the posture or position of the worm. On the other hand, since the texture and outline cannot be used as a clue when parts of the bodies are overlapping in a binarized image, information of preceding and succeeding posture images is always necessary in WormTracer.

Although WormTracer uses several hyperparameters that need to be pre-determined, such as the loss weights, speed (see method), and learning rate of stochastic gradient optimization, experiments on *dpy* mutants and movies with low frame rate have confirmed that, in practice, the recommended parameters can be used to obtain the correct centerlines even in non-canonical cases.

Conventional methods for acquiring the centerlines include those that use superposition of eigenworms and those that use CNN without utilizing temporal continuity. However, they sometimes fail to acquire the correct centerline or to reproduce details in images where the body texture is not recognizable. Our method addresses such issues and enables more accurate centerline acquisition for a larger dataset than existing methods.

This method is optimized based on the assumption that consecutive videos taken over time are used as input data. Therefore, it is not suitable for application to single images or images with extremely low frame rates. However, it was confirmed to work effectively for both high-frequency data, such as 66 fps, and low-frequency data, such as 5 fps, making it applicable to a wide range of frame rates. The limitation of WormTracer will arise when the worm assumes a complex posture that is difficult to analyze, and the interval between shots is so long that the correct centerline cannot be inferred from the posture immediately before or after.

Information on the centerlines of a worm may be used for detail-oriented purposes, such as analysis of head swing movements [19][20][21][22]. When obtaining centerlines for such purposes, even the slightest error in the centerline of the head can have a significant impact. In many cases, the centerlines obtained by existing methods were partially imprecise for the head and tail of the body, even though the overall shape was roughly correct. By using WormTracer, analyses can be performed that focus more on detailed motion than is possible with conventional methods.

Currently used eigenworms are the principal components obtained from the centerlines of the worm assuming simple postures [8]. Therefore, the new eigenworms obtained from the centerlines including the complex postures obtained by WormTracer, are expected to more accurately represent worm postures than the existing eigenworms. In fact, when compared to the existing eigenworms, the new eigenworms more accurately represented the worm postures in the database. For example, there has been active research on the phenomenon of sequential contraction of muscles in a wave-like pattern during forward and backward movement [25][26][27]. When worm locomotion is reduced to a lower dimension using eigenworms in such studies, a new set of eigenworms created based on data that includes complex postures would facilitate a more precise and detailed analyses of the movements, such as those in which the ventral or dorsal side contracts in unison.

## Materials and Methods

### Worm strains used

*C. elegans* worms were cultured at 20°C on NGM plates and *E. coli* strain OP50 was used as food source. Bright field images of adult individuals of the QD15 #4 strain (*Is[H20p::nls3::tdTomato, H20::nls2::GCaMP6f], qjIs11[glr1p::svnls2::tagBFP, ser2(prom2)p::svnls2::tagBFP], Is[eat4p::svnls2::mCardinal, lin44p::GFP*]) , unless otherwise noted, or the *dpy-20(e1370)* IV strain were obtained.

### Image capture and binarization of worms

In this study, we used time-lapse images captured from worms moving freely on a plate covered with a highly transparent gel called Phytagel (Merck). The images were taken at 66 fps and an infrared red camera was used. The binarization threshold of the captured images was determined manually using ImageJ; only the sequence of binarized images was needed for the analysis by WormTracer.

### Centerline determination by a classical method (step 0)

If a worm is in a simple posture, the centerline is easily obtained by applying a thinning algorithm to the binary image. As a preprocessing step in WormTracer, objects other than the largest object in binary images are eliminated and holes are filled. This will eliminate obstacles such as eggs laid by the worm or reflections by impurities or scratches on the gel. Next, the centerline of the worm is obtained by Zhang’s thinning process [9]. If branching exists after thinning, a line connecting the two most distant points along the line by the shortest path is obtained as the centerline (Fig 10). In practice, appropriate centerlines can be obtained in general for simple postures, while correct centerlines are difficult to be obtained for complex postures (Fig 11).

**Fig 10.**
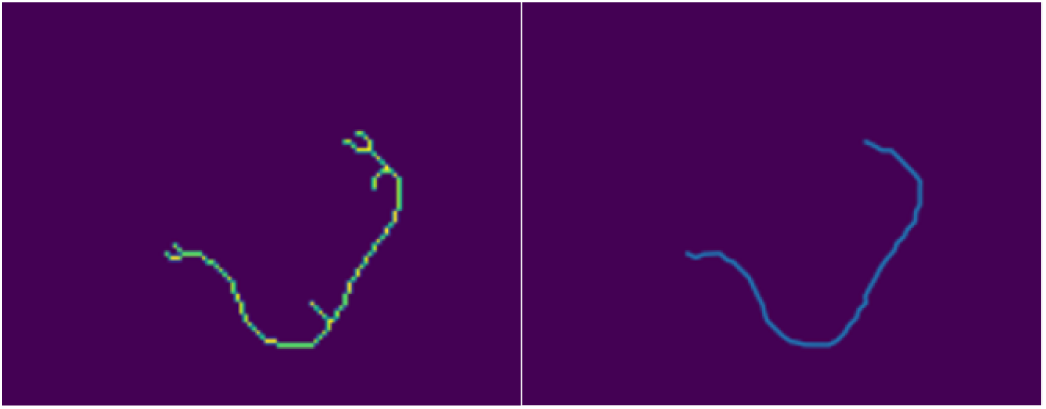
Thinning results. Left: Example of branching caused by thinning Right: Result of selecting the longest path

**Fig 11.**
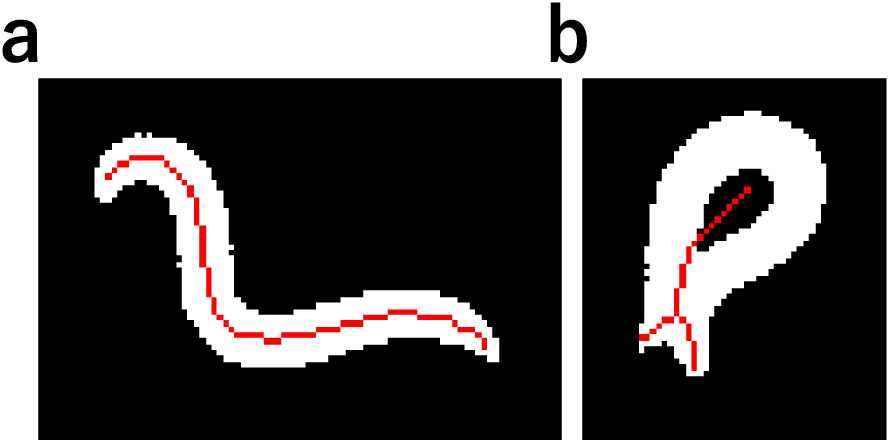
Acquisition of the skeleton by the thinning process. The red line is the result of thinning. a: In a simple posture, the centerline was successfully obtained by the thinning process. b: An example where the centerline was not well obtained by the thinning process in a complex posture.

In the following step, the centerlines are represented by line segments connecting a fixed number of junction points (including end points) so that the distance between adjacent junction points, hence the length of each line segment, *l*, is constant (this value is also called unitLength). The centerlines obtained by thinning were converted to this format by a linear interpolation. Therefore, the posture and position of the worm can be expressed by a set of angles of each line segment, relative to the coordinate axis, and the coordinate of the centroid of the segment junction points (Fig 12).

**Fig 12.**
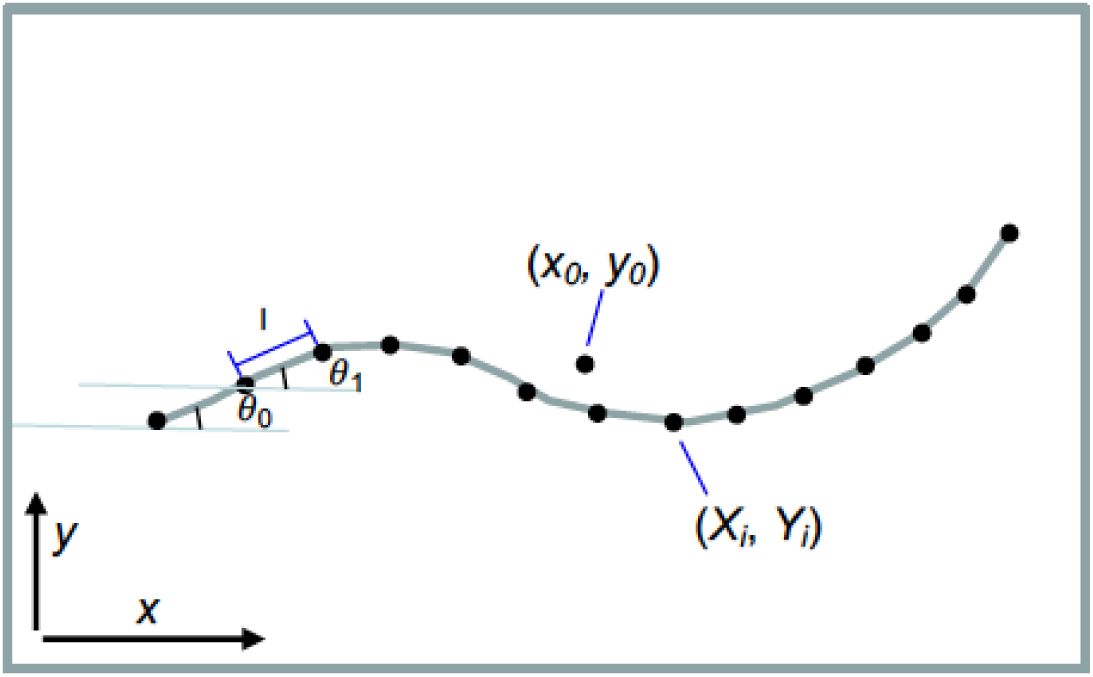
Quantification of centerline. The default number of points that make up the centerline is *M*+1=100. *θ_i_* is the angle between the x-axis and the line segment connecting the i-th point (= (*X_i_* , *Y_i_* ) and the *i* + 1-th point (= (*X_i+1_* , *Y_i+1_*) ). The distance (*l*, unitLength) between adjacent component points is constant. (*x_o_*, *y_0_*) represents the centroid of the points constituting the centerline.

### Model worm and model image generation

To evaluate the estimated centerline, a model worm is generated along the centerline and compared to the real image. The main variable that determines the shape of the model worm is the width *w_i_* as a function of the position (*i*) along the centerline, which is defined as follows

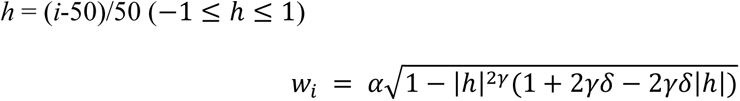

where *α* represents the thickness of the worm, *γ* is the extent of the taper length, and *δ* is the extent of sharpness of the taper (Fig 13). This function is a modification of an ellipse function 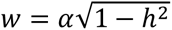 (solution of 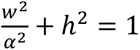) where *h* axis is transformed non-linearly while maintaining continuity of the first derivatives of the resulting function. These three hyperparameters are fixed for the first half of the optimization and added as adjustable shape parameters in the second half of the optimization.

**Fig 13.**
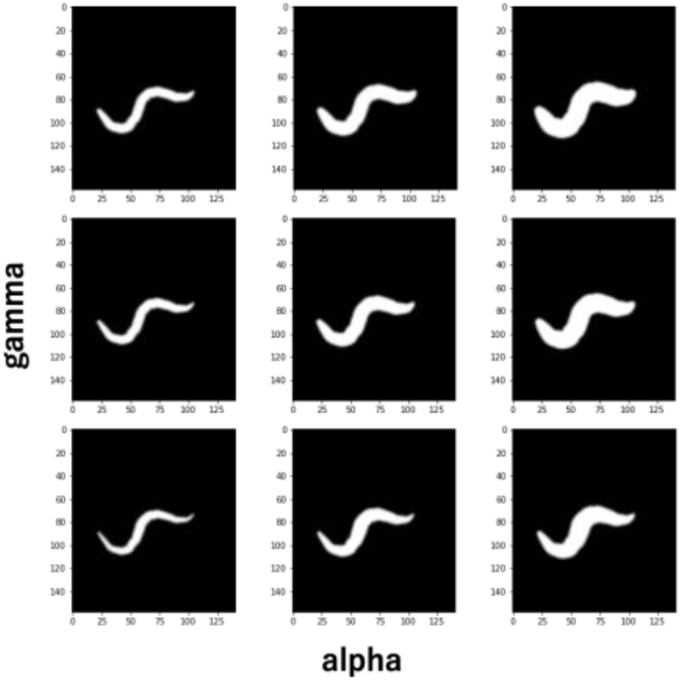
Changes in the model image as α and γ are varied. The larger *α* is, the thicker the worm becomes, and the larger *γ* is, the thicker the worm becomes to the tip.

While the input images are binarized, the luminance values of the model worm images need to be continuous because we adopt gradient-based optimization where the loss function includes image loss. Thus, a model image suitable for optimization is created using a sigmoid function whose luminance value changes smoothly depending on the distance d from the midline, and steeply drops at *w_i_* determined as above.

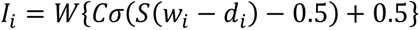

where *I_i_* is the luminance value at a pixel at a perpendicular distance of *d_i_* from the *i*’th centerline segment and *w_i_* is the width of the model worm as defined above, *C* is contrast, *σ* is a sigmoid function. *W* (white_value) is the value of the white portion of the binary image, which is 255 when OpenCV is used.

### Selection of the boundaries of the blocks to be used for optimization

The time series of worm images is divided into several blocks for optimization. There are two reasons for this operation: first, to reduce the amount of memory used at each optimization; second, the centerlines of the complex postures can be estimated using the centerlines of simple posture as a stepping stone, by sandwiching the complex postures with the previously optimized simple postures at both ends of the optimization block (Fig 2).

Since the frames that serve as the boundaries of blocks are used as footholds, it is necessary that the centerlines taken by thinning are accurate at the ends of the blocks. For this purpose, worm reconstruction is performed on the centerlines obtained by thinning (step 0 centerlines) and the reconstructed images are compared to the input images to determine whether the step 0 centerlines have been taken correctly (Fig 14).

**Fig 14.**
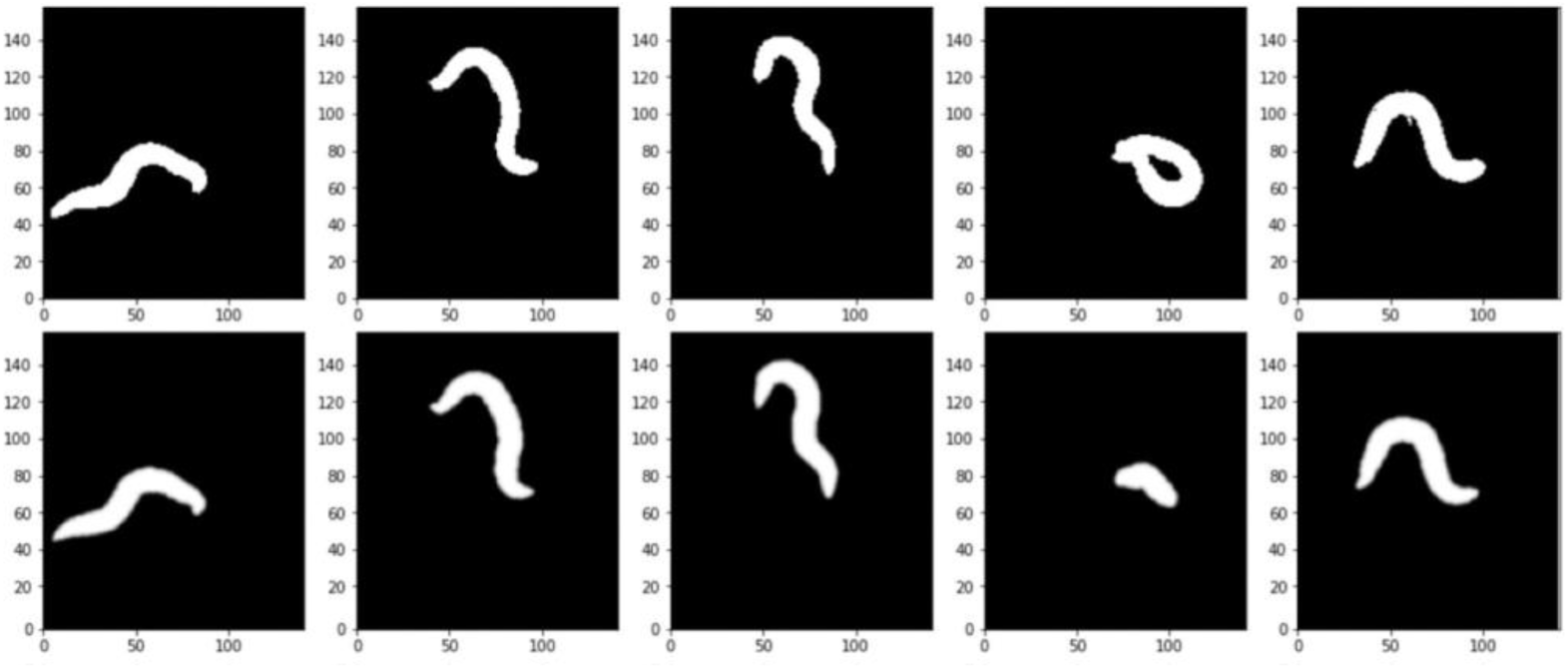
Comparison of the actual binary images (top row) and the image reconstructed from estimated centerlines (bottom row). The second image from the right is not selected as the boundary of the section because it is not well delineated.

The frames with a large difference between the reconstructed worm images and the real images are considered frames when the worm assumes a complex posture. The sequence of images was divided into blocks that enclose such frames (Fig 15).

**Fig 15.**
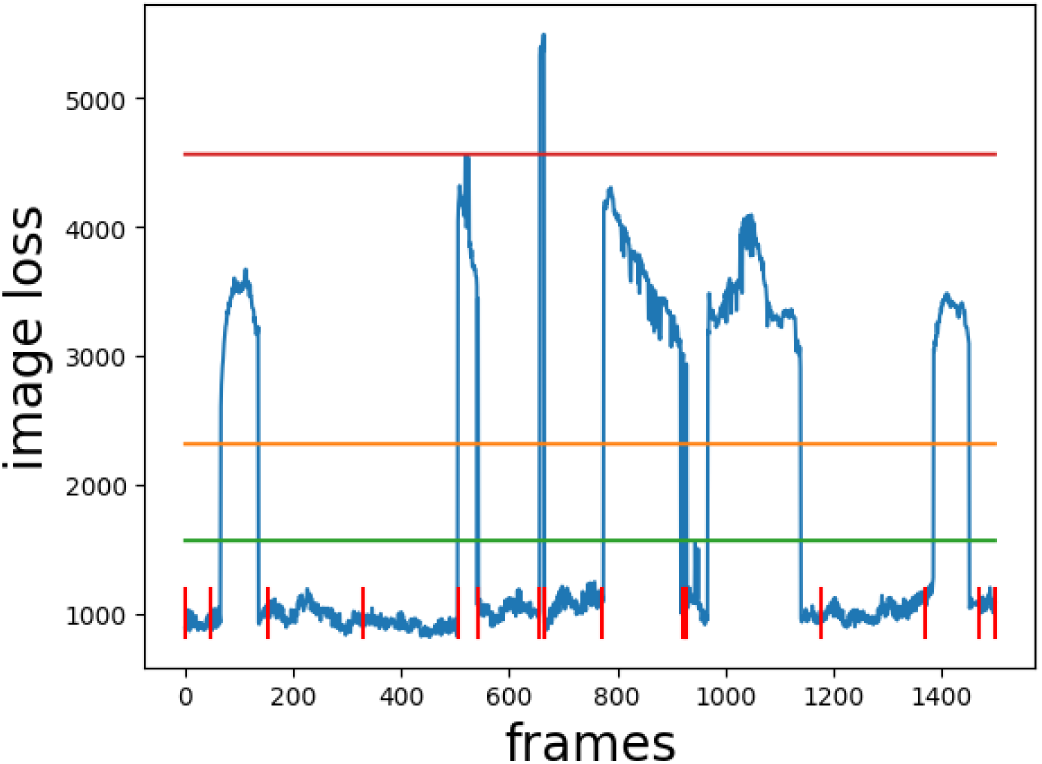
Comparison of the actual binary image and the image reconstructed from the centerline obtained by thinning. The horizontal axis is time and the vertical axis is the mean squared error of all pixels in the images. The red horizontal line represents the theoretical maximum value of squared error; the error relative to the blank image. The orange and green lines are used for detecting complex postured sections; orange represents the 40% value between the red line and the minimum image-loss, and green represents the 20% value. The red vertical bar represents the partition of the blocks used for optimization. The section of the images that are greater than 25% line and some of which is greater than 50% line was considered complex.

### Optimization for each block (steps 1-3)

For each block with the boundaries obtained as above, centerlines are estimated as follows.

In the following description, the beginning and end of the block are depicted as *t* = *t_s_* and *t*=*t_e_*, respectively.

### Optimization parameters and initial values

Pytorch’s gradient-based optimizer, Adam, was used. The process of optimization consists of two steps: a coarse optimization in the first half and a more precise fine-tuning step in the second half. The parameters in the first half to be optimized are the angles of each line segment (*i*’th segment) of the centerline (*θ_t,i_*), the position of the centroid (*x_c,t_*, *y_c,t_*), and *l_t_* (unitLength), for each time frame (*t*). The reason why the length *l_t_* of the worm is included in the optimization parameters is that the length of an actual worm changes slightly due to muscle contraction, so fixing the segment length of the model does not always result in correct centerlines (Fig 16). In the second half of the optimization, shape parameters are also optimized as described later. In the gradient method, the optimization proceeds toward the neighborhood minima of the loss function (described below), so the initial values need to be close enough to the real centerline. Accordingly, the initial values are set as follows.

**Fig 16.**
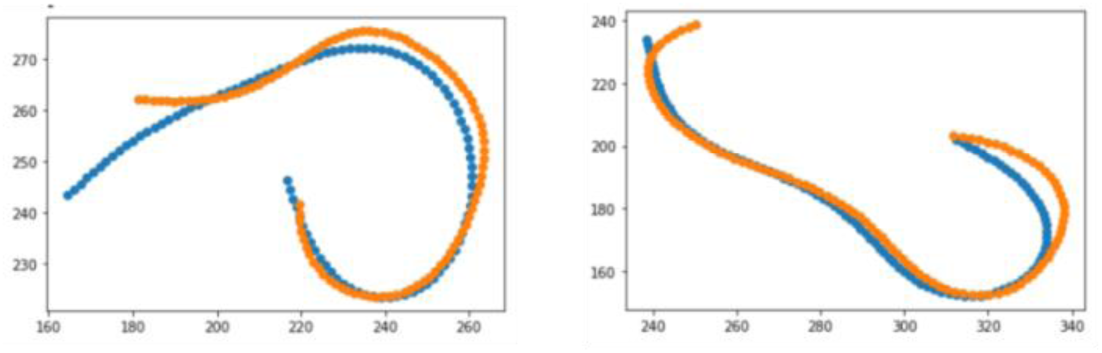
Result of centerline acquisition with fixed unit length. The orange plot shows the centerline obtained by the thinning method, which almost exactly represents the posture of the worm. On the other hand, the blue plot shows the centerline obtained by optimization with constant length, which is longer than the worm in the left image and shorter in the right image. This example indicates that the length of the worm is not constant but varies over time.

- centroid (*x_c,t_*, *y_c,t_*) : The initial values are set to the position of the centroids of the binary images.

- theta (*θ_t,i_*) : The linear interpolation of the *θ* values at the first and last frames of the block (*θ_ts,i_*and *θ_te,i_*) is set as the initial value. *i* = 0, 1, 2,. , *M*-1.

- unitLength (*l_t_*) : The average of the unit lengths between the points constituting the centerline obtained by the thinning, over all frames in the blocks including only simple postures, is set as the initial value. Function of *t*.

Since the model worm is symmetrical, the same worm silhouette is produced with either end of the centerline set as the head (*i*=0). Thus, there are two possible centerline layouts to reconstruct an image that is close to the real worm. Furthermore, the number of rotations of the worm during the complex movement cannot be determined from only the images of the simple postures at ends of the block. Because initial values of *θ* for the whole block is determined by linear interpolation of θ at both ends (*θ_ts_* and *θ_te_*) as described above, the different values of *θ_ts_* and *θ_te_*, even though synonymous for the end images, lead to different initial values. If the number of rotations between the ends are different from the movement of the real worm, it is impossible to optimize correctly using the gradient approach (Fig 17a).

**Fig 17.**
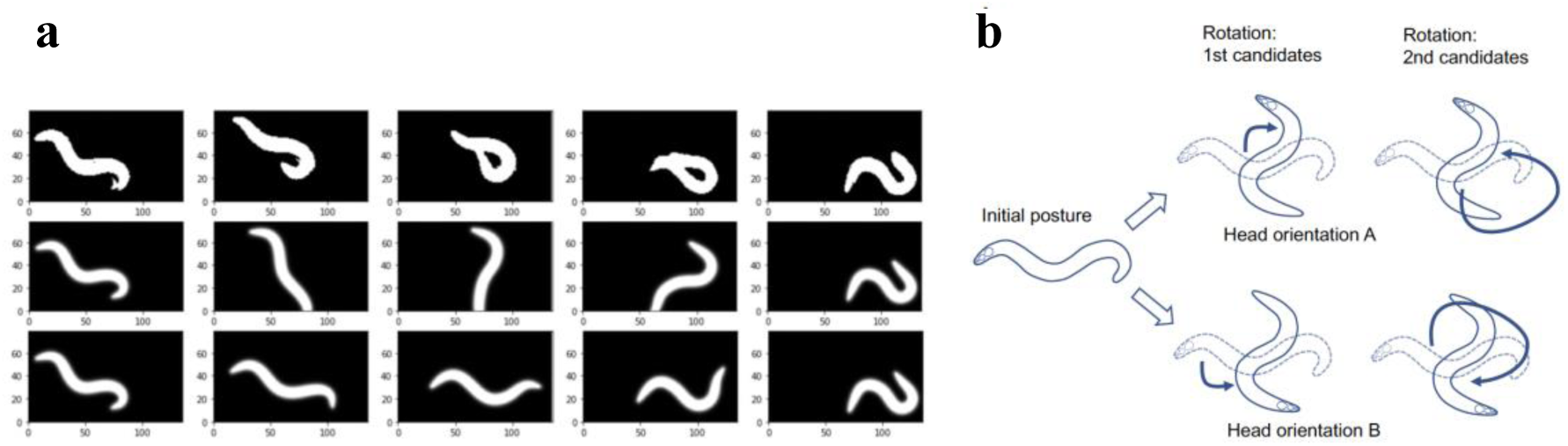
Effect of the choice of initial values of centerlines. a real images and initial images reconstructed from initial values of centerlines. The top row is the binarized image of the raw data. The second row depicts the initial values with the correct head orientation correspondence. The third row depicts the initial values for the incorrect head orientation correspondence. b Schematical representation of possible initial values of rotation angles.

Therefore, first, the head orientation is chosen arbitrarily for the starting time point *t_s_*. For the last time point *t_e_*, two possibilities with either end of the centerline as a head (depicted as the A orientation and the B orientation) are considered (Fig 17b). For each head orientation, among the freedom of +/- 2*nπ* for *θ_te_* (n being an integer), those with smallest rotation from *θ_ts_* are chosen as the first candidates for *θ_te_*. The first candidates are the ones with the smallest MSE (mean squared error) between *θ_ts_* and *θ_te_*, and the second candidate is the one with next smallest MSE, which usually corresponds to the rotation to the opposite direction. As described below, the results of optimization using the first candidates are evaluated, and if both optimization results are judged incorrect, the second candidates are used.

### Posture estimation by optimization

The optimization is performed using the gradient method Adam implemented in Pytorch. The gradient-based optimization is performed for each of steps 1, 2, 3 (Fig. 1), of which steps 2 and 3 are subdivided to substeps A and B, each for orientation A and B (Fig 15b), although step 3 is performed only when step 2 was unsuccessful (see below). The orientation that resulted in the smaller LOSS is chosen.

Each of steps 1, 2A, 2B, 3A and 3B are performed in two phases: in the first half, the approximate midline shape is estimated, and in the second half, the details are fine-tuned. In the first half, the following five types of loss are set: image loss, temporal angular continuity loss, spatial angular continuity loss, length continuity loss, and centroid loss, which are summed up to make the objective LOSS. Each loss is calculated as follows, where *w_im_*, *w_con_*, *w_smo_*, *w_len_* and *w_cen_* are loss weights. *M* is the number of segments (=99 in default). *θ_t,i_* is the angle of i’th centerline segment at time *t* (*i* = 0,1,2,3,…. *M*-1, *t_s_* ≤ *t* ≤ *t_e_*). Image loss: *w_im_* multiplied by MSE between the binary image of the actual worm and the image of the worm reconstructed from the centerline at *t*.

Temporal continuity loss: MSE between the corresponding angles at the previous and next time points. Namely, 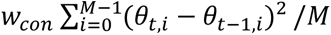

Spatial continuity loss: MSE between adjacent angles at the same time. Namely, 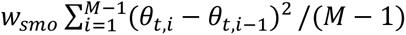

Length continuity loss: the lengths between the segment junction points at the previous and next time points. Namely, *w_len_*(*l_t_* – *l_t_*_–1_)^2^

Centroid loss: square error between the centroid of the actual binary image and the centroid of the centerline at time *t*. Namely, 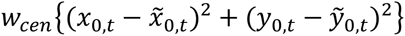 where *x*_0,*t*_ and *y*_0,*t*_ represent coordinates of the centroid of junction points for estimated centerline, while *x̃*_0,*t*_ and *ỹ*_0,*t*_ represent that of the centroid of the binarized image, at time *t*

In performing optimization, an annealing method is used for image loss and spatial continuity loss. By applying an annealing function *F_a_* to the loss weight of a block; *w_im_* = 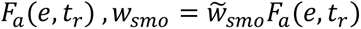 where *e* is epoch number, *t_r_* is the frame position in the block, *w̃_smo_* is basal temporal continuity loss weight. The loss is reflected from both ends of the block as the optimization proceeds (Fig 18). Thus, the centerline is not optimized for the entire block at the same time, but optimization gradually proceeds from the ends of the block where the correct centerline is already estimated. Starting from the images where the correct posture is estimated at both ends, optimization is extended toward the center of the block. Therefore, the accuracy increases as optimization proceeds by utilizing the temporal continuity of the worm’s posture and by utilizing the correct posture as a stepping stone (Fig 19). The rate of change of the loss weight distribution is determined by a hyperparameter called speed.

**Fig 18.**
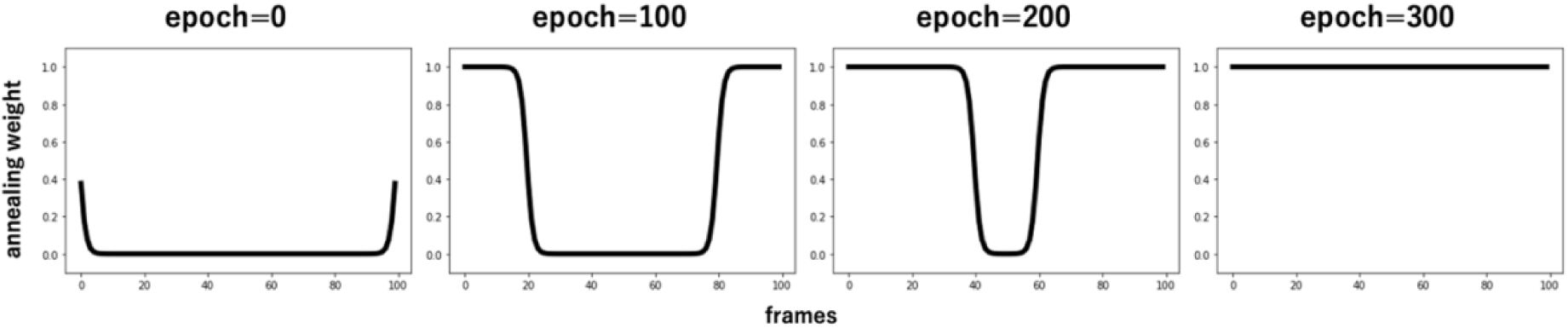
Time variation of weight distribution in annealing. In the early stages of optimization, the frames near both ends have larger loss weights and therefore is optimized, and the optimization proceeds gradually to the center of the optimization block. Note that the speed of optimization progression depends on a hyperparameter called speed.

**Fig 19.**
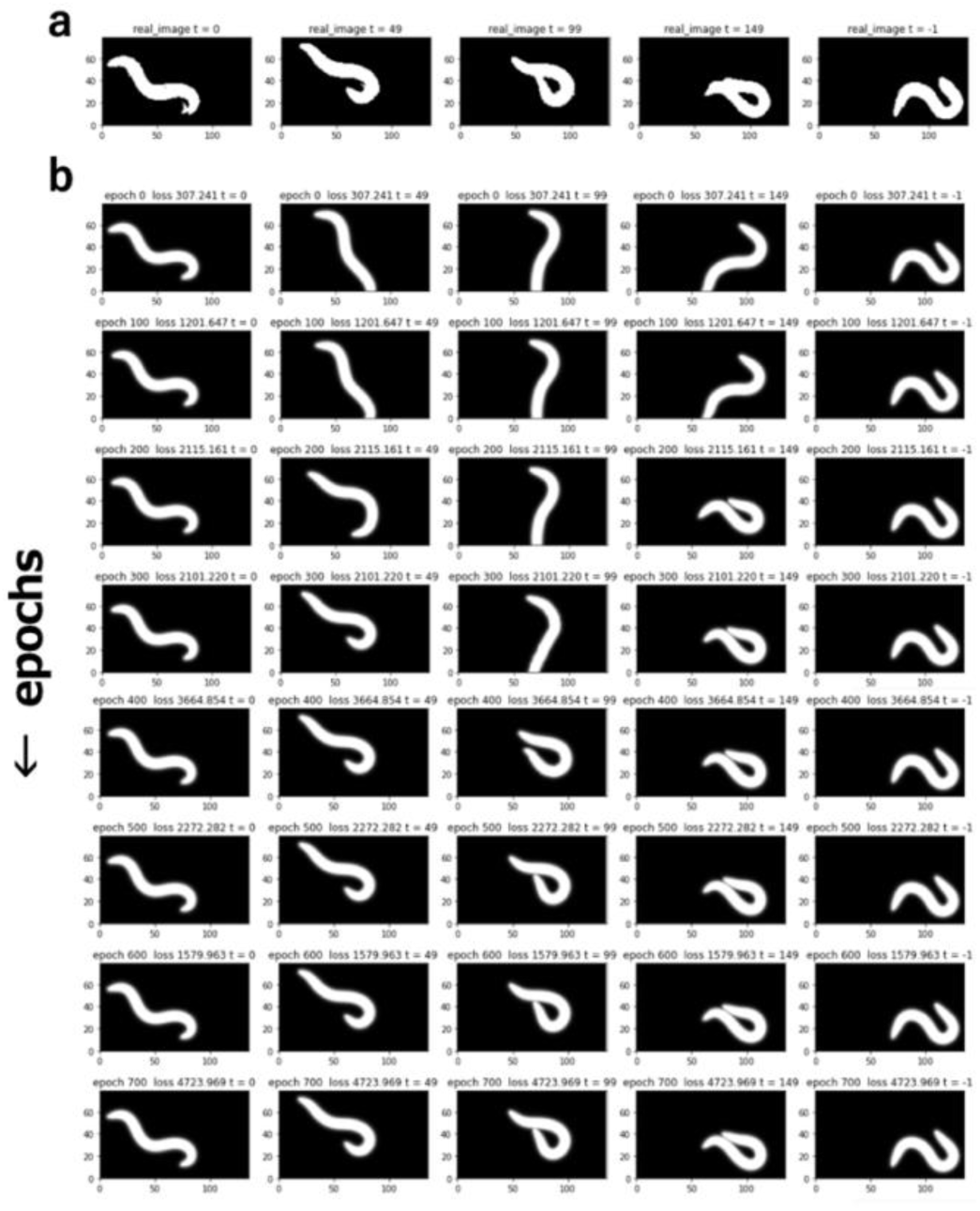
Gradual progression of optimization from the ends of the block when annealing is applied. a Binarized real image. b Optimization progresses from the end frames as the epoch progresses.

In the fine-tuning step, the shape parameters *α*, *γ*, and *δ* are included in optimizable parameters, while centroid loss is disregarded. By performing the optimization again under these conditions, the details of the centerline, especially at the tips of the worms, are fine-tuned (Fig 20).

**Fig 20.**
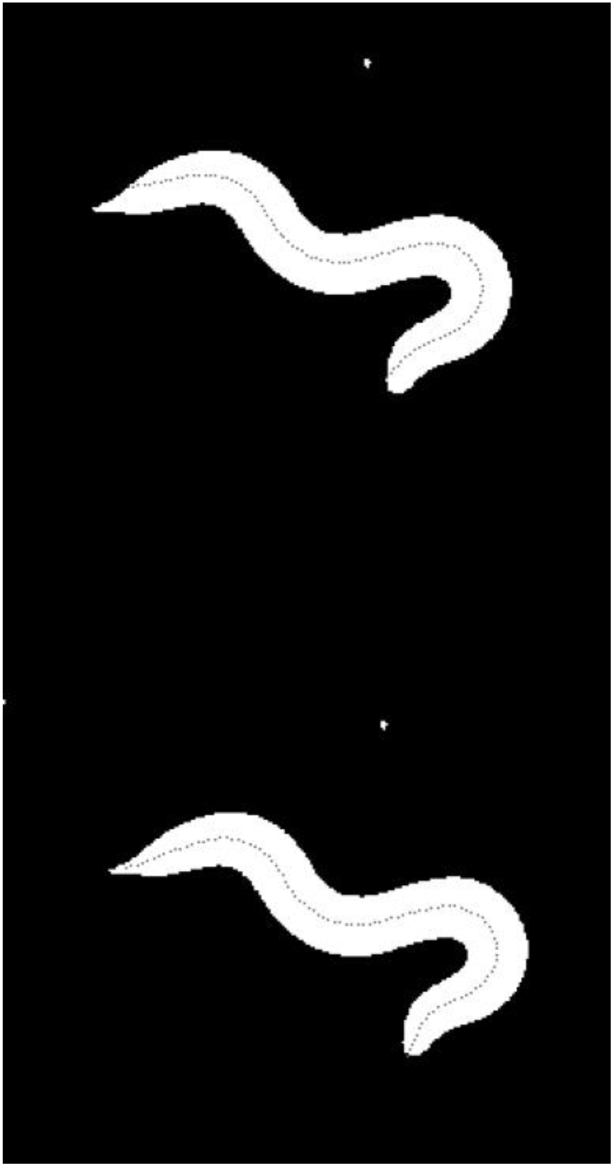
Change of the centerline before and after fine-tuning. Top: Centerline obtained in the first half of the optimization step Bottom: Fine-tuned centerline in the second half of the optimization step

### Detection of optimization failure

After the step 2 optimization, attained loss is evaluated for each block. Blocks that had a significantly large loss, where the difference between the maximal LOSS values in the block and median LOSS value of all frames is greater than four times the quartile range of all frames, are considered failure and centerline is estimated again (step 3) using a different set of initial values of *θ*. As described earlier, as the initial values of *θ*, those with the second smallest rotation angles are used (Fig 17b, right). The results are then updated if the loss is smaller than the step 2 results.

### Head and tail flipping detection and correction

Considering that head-tail exchange gives the same image in our model worm, the overall centerlines obtained may have inverted ends at the junctions of the optimization blocks. Therefore, in order to check whether there is an abrupt change of centroid direction in the previous and next frames, the MSE between the coordinates of segment junction points for adjacent frames at block junctions is calculated by using the estimated centerline and those from an inverted centerline. If the MSE is smaller with the inverted coordinates, the inverted centerlines are adopted for one of the blocks.

### Head Determination

It is known that worms generally swing their heads more frequently than their tails. Therefore, the 99 angles between adjacent centerline segments (*θ_t,i+1_*-*θ_t,i_*) are calculated, and the end with the larger variance of curvature is recognized as the head throughout all frames, and the order of angles is modified to start from the head (Fig 21).

**Fig 21.**
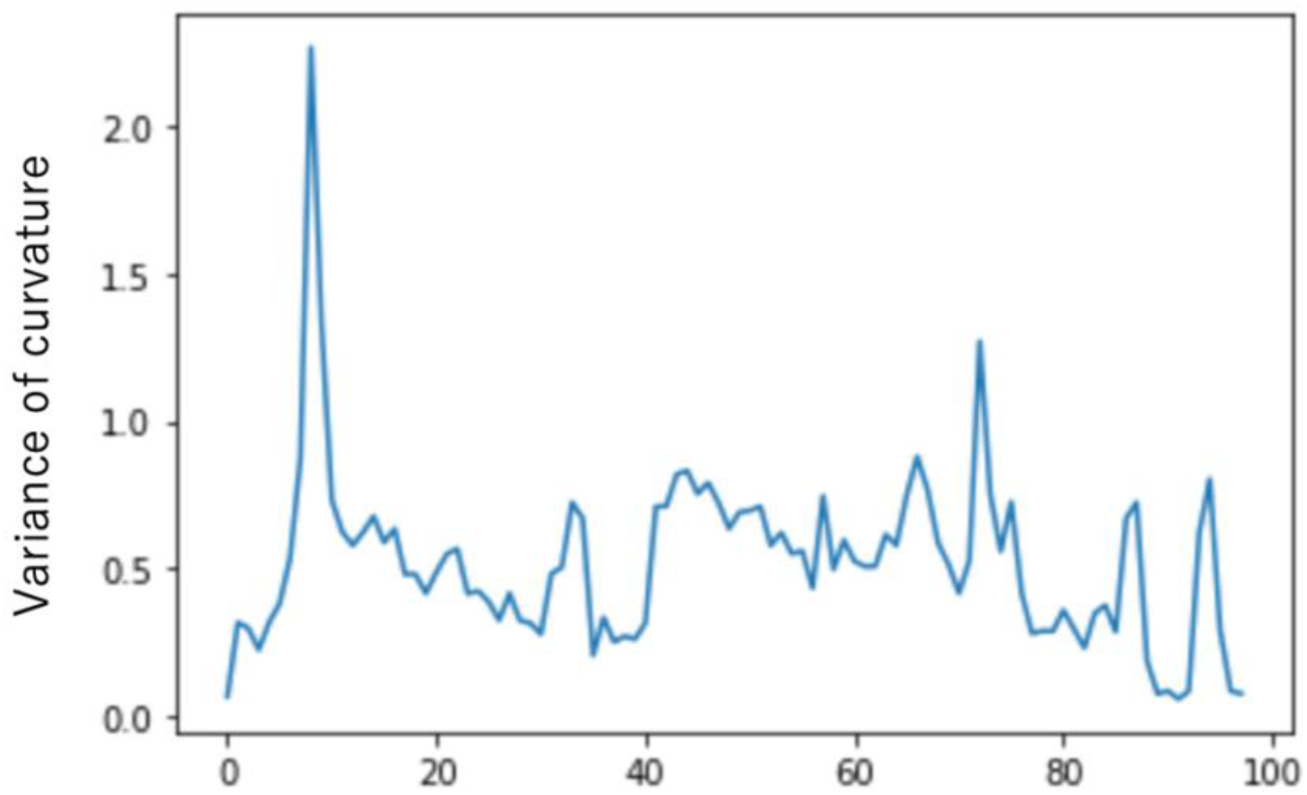
Determination of head based on variance of curvature. Horizontal axis shows the number of body segments, while vertical axis shows variance of curvature along time in optimized centerlines. The curvature variance on the side with the smallest index is large, and this side is judged to be the head.

### Comparison with existing methods

To compare WormTracer with existing methods, centerlines were estimated for the same image sets by using WormPose and EigenWormTracker in addition to WormTracer. To maintain fairness, we used the sample images for each method and applied all three methods on each image set. For optimization hyperparameters, all default settings were used (for WT, continuity_loss_weight=10000, smoothness_loss_weight=100000, length_loss_weight=50, center_loss_weight=50, loss_weight=50, with necessary settings such as Worm_is_lighter in WP modified as instructed).

In order to evaluate the three methods, ground truth was prepared manually. In order to minimize individual bias, handwritten centerlines created by two volunteers were averaged and used. For comparison, the squared error between segment angles (*θ*) was calculated and then averaged for 99 segments within the same time frame.

## Data and Code Availability

All data and code used for centerline determination from worm images are available on a GitHub repository at https://github.com/yuichiiino1/WormTracer.

## Acknowledgments

This work was supported by the CREST program “Creation of Fundamental Technologies for Understanding and Control of Biosystem Dynamics” (JPMJCR12W1) of the Japan Science and Technology Agency (JST), and Grant-in-Aid for Scientific Research (JP17H06113, JP22H00416, 20K21805, JP25115009 and 19H04980). YT was supported by JST PRESTO (JPMJPR1947) and Grants-in-Aid for Scientific Research (JP26830006, JP18K14848, JP16H01418, JP18H04728, JP17H05970 and 19H04928). We are grateful to Ibuki Ohmori and Moe Hosomi for creating handwritten centerlines used in this work.

## Notes

### Competing Interest Statement

The authors have declared no competing interest.

